# Partitiviruses infecting *Drosophila melanogaster* and *Aedes aegypti* exhibit efficient biparental vertical transmission

**DOI:** 10.1101/2020.06.01.128819

**Authors:** Shaun T. Cross, Bernadette L. Maertens, Tillie J. Dunham, Case P. Rodgers, Ali L. Brehm, Megan R. Miller, Alissa M. Williams, Brian D. Foy, Mark D. Stenglein

**Affiliations:** Department of Microbiology, Immunology, and Pathology, College of Veterinary Medicine and Biomedical Sciences, Colorado State University, Fort Collins, CO, USA; Department of Biology, Colorado State University, Fort Collins, CO, USA

**Keywords:** *Drosophila melanogaster*, *Aedes aegypti*, galbut virus, verdadero virus, partitivirus, arthropod, vertical transmission, metagenomics, virus discovery, virome

## Abstract

Partitiviruses are segmented, multipartite dsRNA viruses that until recently were only known to infect fungi, plants, and protozoans. Metagenomic surveys have revealed that partitivirus-like sequences are also commonly associated with arthropods. One arthropod-associated partitivirus, galbut virus, is extraordinarily common in wild populations of *Drosophila melanogaster* fruit flies. To begin to understand the processes that underlie this virus’s high global prevalence, we established colonies of wild-caught infected flies. Infection remained at stably high levels over three years, with between 63-100% of individual flies infected. Galbut virus infects fly cells and replicates in tissues throughout infected adults, including reproductive tissues and the gut epithelium. We detected no evidence of horizontal transmission via ingestion but vertical transmission from either infected females or infected males was ~100% efficient. Vertical transmission of a related partitivirus, verdadero virus, that we discovered in a laboratory colony of *Aedes aegypti* mosquitoes was similarly efficient. This suggests that efficient biparental vertical transmission may be a feature of at least a subset of insect-infecting partitiviruses. To study the impact of galbut virus infection free from the confounding effect of other viruses, we generated an inbred line of flies with galbut virus as the only detectable virus infection. We were able to transmit infection experimentally via microinjection of homogenate from these galbut-only flies. This sets the stage for experiments to understand the biological impact and possible utility of partitiviruses infecting important model organisms and disease vectors.

**Importance:** Galbut virus is a recently discovered partitivirus that is extraordinarly common in wild populations of the model organism *Drosophila melanogaster*. Like most viruses discovered through metagenomics, most of the basic biological questions about this virus remain unanswered. We found that galbut virus, along with a closely related partitivirus found in *Aedes aegypti* mosquitoes, is transmitted from infected females or males to offspring with ~100% efficiency and can be maintained in laboratory colonies over years. This represents one of the most efficient means of virus transmission described, and likely underlies the successful spread of these viruses through insect populations. We created *Drosophila* lines that contained galbut virus as the only virus infection and showed that these flies can be used as a source for experimental infections. This provides insight into how arthropod-infecting partitiviruses may be maintained in nature and sets the stage for exploration of their biology and potential utility.

## Introduction

Metagenomic surveys of wild organisms have revealed a breathtaking abundance and diversity of viruses (1–10). Some recent studies have described hundreds or thousands of new virus or virus-like sequences (1, 2, 4, 10). These have contributed substantially to our understanding of virus evolution and genome structure, have expanded the known host range of some virus types, and have led to the establishment of entirely new families of viruses (11).

The explosion of virus discovery from metagenomics is an important advance in virology, but questions remain about the biological impact of these viruses. Virus sequences can often not even be confidently assigned to a particular host, since many metagenomic datasets derive from intact organisms (holobionts) or from pools of organisms. A virus sequence could therefore represent infection of microbiota of the targeted organism. For example, Webster *et al.* described a variety of novel virus-like sequences associated with wild *Drosophila* fruit flies (12). A subsequent study on one of these viruses, Twyford virus/Entomophthovirus, revealed that it was in fact a virus of a fungal parasite of flies (13).

Like many groups of RNA viruses, the partitiviruses (family *Partitiviridae*) have undergone a recent expansion via metagenomics (2, 12, 14–18). Partitiviruses were previously only known to infect plants, fungi, and protozoa, and what is known about these viruses is from studies in these hosts (19–24). Partitiviruses genomes are composed of two or more segments of double stranded (ds) RNA. The particles of these viruses are non-enveloped and multipartite: individual particles contain a single segment. Although infections are persistent and generally not associated with clear phenotypic differences, partitivirus infection can alter hosts in measurable ways. Partitivirus RNA levels were correlated with increased fecundity of *Cryptosporidium parvum* parasites (25). Some partitiviruses of fungal pathogens confer hypovirulence to their hosts (26–29). A partitivirus of jalapeño pepper plants made infected plants less attractive to aphids (30).

We had previously observed partitivirus-like sequences in wild-caught *Anopheles gambiae* mosquitoes from Liberia, Senegal, and Burkina Faso (14). Partitivirus-like sequences have also been identified in a variety of other types of mosquitoes (8, 14, 17, 18, 31). At the time we identified the partitivirus-like sequences in *Anopheles* mosquitoes, among the most closely related sequences were those of galbut virus, a partitivirus-like sequence that had first been identified in *Drosophila melanogaster* fruit flies (12). Galbut virus was found to be literally ubiquitous in sampled populations of *D. melanogaster* and present in related species in the melanogaster group (12, 16, 32, 33). Although small RNA profiles suggested that galbut virus legitimately infected fly cells, it was not clear whether this was indeed the case. Recognizing this uncertainty, Webster et al named galbut after the Lithuanian word galbūt, meaning maybe (12).

We therefore undertook studies to better understand the biology of partitiviruses that were common constituents of the viromes of important model organisms and disease vectors. We established colonies of wild-caught galbut virus infected *D. melanogaster*. We also discovered that a laboratory colony of *Aedes aegypti* mosquitoes harbored a pre-existing infection of a related partitivirus, which we named verdadero virus. We confirmed infection in *Drosophila* cells, quantified efficiency of horizontal and vertical transmission, isolated galbut virus through inbreeding, and established a system for experimental infection by microinjection.

## Materials and Methods

### *Drosophila* Collections

Wild *Drosophila* were collected in Fort Collins, Colorado, USA, from a backyard compost bin. Subsets of flies were moved to rearing bottles for colony establishment or stored at −80°C. Laboratory reared *D. melanogaster* strain w^*1118*^ were provided by Dr. Susan Tsunoda. Additional stocks of flies from the Drosophila Genetic Reference Panel (DGRP; strains 360, 399, and 517) were obtained via the Bloomington Drosophila Stock Center (34).

### Arthropod Maintenance and Rearing

Both wild-derived and laboratory-derived *Drosophila* were reared at 25°C on the Bloomington Drosophila Stock Center Cornmeal *Drosophila* medium (https://bdsc.indiana.edu/information/recipes/bloomfood.html). *Drosophila* stocks were moved to new bottles every 2 weeks.

*Aedes aegypti* strain Poza Rica (partitivirus infected) (35) and strains New Orleans and Vergel (partitivirus uninfected) (36) were used for vertical transmission experiments. Mosquitoes were reared as previously described (37).

### Vertical and Horizontal Transmission Experiments

To quantify vertical transmission of galbut virus, virgin male and female flies were crossed. Infected wild-caught colonized flies (colony FoCo-17) and uninfected flies (strain w^*1118*^) were used for experiments. Individual mating pairs were maintained in the same bottle with an apple agar egg plate and yeast paste (1:1 yeast and water) to promote egg laying. Egg plates were removed daily and eggs collected. Eggs were placed in a 50% bleach solution for 2 min to remove the outer chorion layer, after which they were immediately transferred to clean ddH2O to remove residual bleach. Bleached eggs were then placed either in vials containing media either together (experiment 1) or as individuals (experiment 2), allowing offspring to rear to adulthood. FoCo-17 parents from which the eggs were derived were tested via quantitative reverse transcription polymerase chain reaction (qRT-PCR) for galbut virus, and if positive, 2-5 day old offspring were collected and also tested via qRT-PCR. Primer sequences were: galbut virus: CCGTGAAGCAAGGAATCAAT, TGCCGATTTTCTGCTCTTTT; RpL-32: TGCTAAGCTGTCGCACAAATGG, TGCGCTTGTTCGATCCGTAAC. Crosses consisted of either a FoCo-17 female and w^1118^ male (maternal transmission) or w^1118^ female and FoCo-17 male (paternal transmission). Exact binomial 95% confidence intervals were calculated with the binom R package.

For vertical transmission of verdadero virus, virgin male and female *Aedes aegypti* were crossed. Verdadero-infected mosquitoes from the Poza Rica colony were crossed with the uninfected New Orleans and Vergel strains. Previous qRT-PCR screening showed that 100% of the males and females of the Poza Rica colony were infected by verdadero virus, while verdadero infection was undetectable in the other colonies. Primer sequences were: verdadero virus: ATATGGGTCGTGTCGAAAGC, CACCCCGAAATTTTCTTCAA. Groups of 30 male and 30 female mosquitoes were placed together for 2 days to allow mating. After this period, female mosquitoes were blood fed with defibrinated calf blood (Colorado Serum Company, Denver, USA) for egg production, and eggs were collected 3 days post blood meal. Eggs were then reared to adulthood and adults collected 0-2 days post eclosion. These offspring were then tested for virus presence via qRT-PCR. Groups consisted of either infected females and uninfected males (maternal transmission) or infected males and uninfected females (paternal transmission). Exact (Clopper-Pearson) 95% binomial confidence intervals were calculated with the binom R package (38).

Horizontal transmission of galbut virus was measured by allowing uninfected (DGRP 399 and 517) flies to ingest homogenate from FoCo-17 line #30 galbut virus infected flies. This homogenate was infectious by microinjection (see below). Homogenate was created by grinding 200 *Drosophila* with a Dounce homogenizer in 150 μL 1X Phosphate-buffered solution (PBS; 137 mM NaCl, 2.7 mM KCl, 10 mM Na_2_HPO_4_, 1.8 mM KH_2_PO_4_). To remove cellular debris, homogenate was spun down at 12,000xg for 1 min, and supernatant was transferred to a new vial. This was repeated for a total of three spins. The 150 μL of homogenate was mixed with 850 μL of 5% sucrose and 5% yeast extract in ddH_2_O. As a negative control, a second food solution was prepared with sterile 1X PBS instead of homogenate. A single drop of blue food coloring was added to the solution to allow visualization of ingestion. Capillary tubes were filled with 7 μL of food solution. Capillary feeder systems were set up following the protocol by Zer et al. (39). Two exceptions to this setup included the lack of piercing holes in the vial and wetting the cap. 3-5 day old virgin female and male flies were starved for 9 hours, after which they were placed in the capillary feeding vials containing either virus inoculated food or control inoculated food. Flies were allowed to feed for 17 hours. Flies that had a blue abdomen were moved to vials containing standard fly medium. Flies were moved to fresh media as needed until 21 days post inoculation, at which point they were frozen at −80°C. Flies were tested for virus presence via qRT-PCR as above.

### RNA Extractions

We developed and validated a high throughput, magnetic bead based method to extract RNA from flies and mosquitoes. Individual flies or mosquitoes were added to a 96-well round bottom plate (Corning catalog #3958) with 1 metal BB ball and 100 μL lysis buffer (5M guanidine thiocyanate, 0.1M Tris-HCl, pH 7.5, 0.01M Na_2_EDTA, pH 8.0, and 6.25mL 2-mercaptoethanol (βME)) and homogenized at 30Hz for 3 min in a TissueLyzer II (Qiagen). 60 μL of 100% isopropanol was added to each tube and incubated for 1 min. To remove cellular debris, samples were spun down in a centrifuge at 1,200xg for 1 min. Supernatant was removed and added and mixed well by pipetting to 96-well plates containing 90 μL RNA magnetic beads (1 mM trisodium citrate, 2.5 M NaCl, 20% PEG 8000, 0.05% Tween 20, pH 6.4, and 1 mL Sera-Mag SpeedBeads (Thermo Scientific) in a total volume of 50 mL) and 10 μL enhancer (Proteinase K 200ug/mL, 20% glycerol, and 0.5% SDS). Samples were incubated in beads for 5-10 minutes. Using magnetic racks, beads were separated from the supernatant by incubation for 2-3 min, and the supernatant was removed. Tubes were removed from the rack, and the beads were washed with 150 μL wash buffer 1 (20% ethanol, 900 mM guanidine thiocyanate, and 10mM Tris-HCl, pH 7.5) for 2 min. Beads were pelleted again using the magnetic rack and supernatant removed. Beads were then washed with 150 μL wash buffer 2 (WB2; 1X Tris-EDTA buffer (10 mM Tris HCl, 1.0 mM EDTA) pH 8.0, and 80% ethanol) for 2 min. Supernatant was removed. Following this step, beads were resuspended in 30 μL of a DNase I mixture, consisting of 3 μL DNase I (NEB), 3 μL 10xDNase I Buffer (NEB), and 24 μL WB2. Beads were incubated in this mixture for 30 minutes at room temperature. Following incubation, tubes were removed from the magnetic rack and resuspended by pipetting with 100 μL binding buffer (5M Gu-HCl, and 30% isopropanol), and incubated for 5 min. Beads were added to magnetic rack and supernatant removed as previously described. Beads were then washed twice with WB2 as previously described. Following the second wash, tubes were left on the magnetic rack and beads air dried for 3 min. To elute RNA from the beads, tubes were removed from the magnetic rack, beads resuspended by pipetting with 25 μL of nuclease-free H2O, and incubated for 5 min at room temperature. Tubes were placed back on the magnetic rack, and 23 μL of the supernatant, containing the extracted RNA, removed. RNA was quantified fluorometrically using a Qubit instrument (Thermo Fisher) and stored at −80°C.

### cDNA Synthesis and Virus Screening

*Drosophila* were screened for various viruses, including galbut virus, via qRT-PCR. First, RNA was subjected to cDNA synthesis: 5.5 μL of RNA was added to 200 pmol of random 15-mer oligonucleotides and incubated for 5 min at 65 °C, then set on ice for 1 min. A RT mixture containing the following was added (10 μL reaction volume): 1× SuperScript III (SSIII) FS reaction buffer (Thermo Fisher), 5 mM dithiothreitol (Invitrogen), 1 mM each deoxynucleotide triphosphates (dNTPs; NEB), and 100 U SSIII reverse transcriptase enzyme (Thermo Fisher), then incubated at 42 °C for 30 min, 50 °C for 30 min, then at 70 °C for 15 min. 90 μL of nuclease-free H_2_O was added to dilute the cDNA to a final volume of 100 μL.

Following cDNA synthesis, qPCR reactions were set up using Luna Universal qPCR Master Mix (NEB) following the manufacturer's protocol. The qPCR reaction was performed on LightCycler 480 (Roche) with the following protocol: 95°C for 3 min, 40 cycles of 95°C for 10s, then 60°C for 45s, and then followed by a melting curve analysis of 95°C for 5s, 65°C for 1 min, and an acquisition starting from 97°C with a ramp rate of 0.11 °C/s and 5 acquisitions per degree. Some qPCR products were validated by running on a 2% agarose gel and Sanger sequencing. Gels were stained with ethidium bromide and imaged on a Gel Doc (Bio-Rad). Primer sequences were: vera virus: CGTCGGGTGTTTAGAGGTAA, TAACGATGGTGTTCCAAGGT; La Jolla virus: ACCGTATGGCGTCGTACTTC, AAAGTATCAGCAGCGCGAAT; Thika virus: CAGCAGGTCCCTTGCTAAAG, TGGTCAGCATATGACCGAAA; Nora virus: GCACCTGGTCGATTGAATCC, CGTTCAGGGCATAGTCAAGC.

### Shotgun Metagenomic Library Preparation

Library preparation for the *Drosophila* samples utilized a KAPA HyperPrep kit following the manufacturer’s protocol. All libraries were sequenced on an Illumina NextSeq 500 instrument and NextSeq 500 Mid Output v2 Kits (Illumina). *w^1118^ Drosophila* samples were made with a final target library size of 380-430bp and were sequenced using paired-end 2×150bp. FoCo-17 and FoCo-18 flies were sequenced using single-end 1×150bp reads.

### Sequence and Data Analysis

Metagenomic sequencing datasets were processed to taxonomically assign viral reads as previously described (40). Species were assigned to individual flies or pools of flies using a competitive mapping approach. A collection of cytochrome C oxidase subunit 1 (COX1) sequences were collected and curated. Sequences were retrieved from the NCBI nucleotide database by BLASTN searching using as a query the *D. melanogaster* COX1 sequence (NC_024511.2:1474-3009) with an e-value cutoff of 10^−12^ (41). Sequences longer than 1400 bp were retained and collapsed into a set of representative COX1 sequences using cd-hit-est and a similarity threshold of 97% (-c 0.97) (42). These operations produced a set of 233 representative COX1 sequences. Quality and adapter trimmed reads from sequencing datasets were aligned to these sequences using bowtie2 with parameters --local and --score-min C,120,1, and the number of reads mapping to the various COX1 sequences were tabulated (43). Code and the set of representative sequences available at https://github.com/scross92/partitivirus_transmission_paper. All additional data analysis scripts can be found at this location as well. All sequencing datasets have been deposited in the NCBI Sequence Read Archive (SRA) under the BioProject accession PRJNA635623. Assembled genome sequences are deposited in GenBank under accessions <GenBank accessions pending>.

### Phylogenetic analysis

Sequences similar to the predicted galbut virus RNA dependent RNA polymerase were retrieved from the NCBI protein database using blastp with an E-value cutoff of 10^−30^. Sequences longer than 400 amino acids were retained and collapsed into a set of representative sequences using cd-hit and a similarity threshold of 95% (-c 0.95). Sequences were aligned using the MAFFT aligner using the --auto setting (44). Multiple sequence alignments were trimmed with the trimAL tool using setting-automated1 to remove uninformative columns (45). The highest scoring model for tree inference (LG+I+G4) was selected using the modeltest-ng software (46). Maximum likelihood trees were inferred using raxml-ng and bootstraps were allowed to run until convergence (47). Trees were visualized using the ggtree R package (48). Code and alignments are available at the above mentioned github repository.

### Antibody creation

Polyclonal rabbit sera targeting the putative galbut virus capsid protein (encoded on RNA 2) was generated by Pacific Immunology (Ramona, California) using the synthetic peptide Cys-QPRRMIRDKPSLREEAHES.

### Western blotting

Whole flies were homogenized in a cold protein extraction buffer (49) containing 20 mM HEPES (pH 7.5), 100 mM KCl, 5% glycerol, 10 mM EDTA, 0.1% Triton X-100, 1 mM DTT and Complete protease inhibitor cocktail (Sigma). Samples were spun for 5 minutes at 12,000 × g and the supernatant was retained. The protein samples were quantified using the BCA Protein Assay Kit (Thermo Fisher) and diluted 1:10. 15 μg of protein were suspended in NuPAGE LDS Sample Buffer (4X) (Thermo Fisher), heated at 70°C for 10 minutes, then loaded onto a NuPAGE 4-12% Bis-Tris, 1.0mm × 12 well gel (Thermo Fisher). Protein samples were separated via SDS-PAGE and transferred onto a 0.45 μm nitrocellulose membrane (BioRad). The membrane was incubated with anti galbut virus capsid rabbit serum at a dilution of 1:100 and pre-immune rabbit serum at a dilution of 1:100. Primary antibody was detected using a fluorescently-labeled goat anti-rabbit secondary antibody (Li-Cor). Blots were imaged using an Odyssey scanner (Li-Cor).

### Immunofluorescence Assay

Slides were prepared according to Stenglein et al. (50). Whole adult *D. melanogaster* tissue sections were prepared for immunofluorescence assays. Adult *Drosophila* were collected, knocked down at 4°C, and legs and wings were removed. Flies were placed in 4% paraformaldehyde in 1X PBS and fixed for 24 hours at 4°C. Following fixation, *Drosophila* were removed from paraformaldehyde and stored in 1X PBS. Fixed *Drosophila* were sent to Colorado State University Veterinary Diagnostic Laboratories’ histology lab for paraffin embedding and sectioning. Sections were deparaffinized in xylene, rehydrated in graded ethanol, placed in 1 mM EDTA at 99°C for 20 min, and rinsed with water. Sections were washed 2 × 5 min in 1X Tris-buffered saline (TBS; 50 mM Tris-Cl pH 7.5, 150 mM NaCl) + 0.025% Tween-20, permeabilized in 1X PBS + 0.1% Triton X-100 for 10 min, and washed 3 × 5 min in TBS + 0.05% Tween-20 (TBS-T). Slides were then blocked with 1X TBS + 1% BSA for 20 min, incubated for 30 min with anti-galbut rabbit serum at a dilution of 1:500 and pre-immune rabbit serum at a dilution of 1:500, washed in TBS-T, incubated for 30 min with Alexa-Fluor 488-conjugated goat anti-rabbit secondary antibody, and washed again. The second wash included 0.5 μg/ml Hoechst 33342. Sections were mounted in Vectashield, coverslipped and sealed with clear nail polish, then imaged with an Olympus IX81 Inverted Confocal Laser Scanning Microscope.

### Intrathoracic Microinjections

100 adult flies from the galbut-only line (FoCo17 line #30) were placed in 200 μL 1X PBS and homogenized by hand with a 1.5 mL homogenizer. Homogenate was spun down at 12,000xg in a microcentrifuge for 1 min to remove cellular debris. This was performed three times to remove all cellular debris. Homogenate was filtered at 0.22 μM filter to remove any cellular material. 50 nL of the homogenate was injected into flies intrathoracically using a Drummond Nanoject II.

## Results

### Galbut virus in local wild *Drosophila melanogaster* populations and establishment of colonies of galbut virus infected flies

Webster et al described galbut virus as an apparently ubiquitous infection of wild *D. melanogaster*. Motivated to better understand the biological impact of galbut virus on infected flies and the dynamics that produced this high global prevalence, we sampled local fly populations and established colonies of wild-caught infected flies. We collected flies from Fort Collins, Colorado and performed metagenomic sequencing to characterize their viromes. Local populations of *D. melanogaster* were as expected infected with galbut virus (**Fig. 1**). We collected flies from the same backyard compost bin over 3 years and found 94%, 84%, and 67% of individual flies infected in 2017, 2018, and 2019 (**Fig. 1A**). We used subsets of collected flies to establish colonies, which we called FoCo-17, −18, and −19.

**Figure 1:**
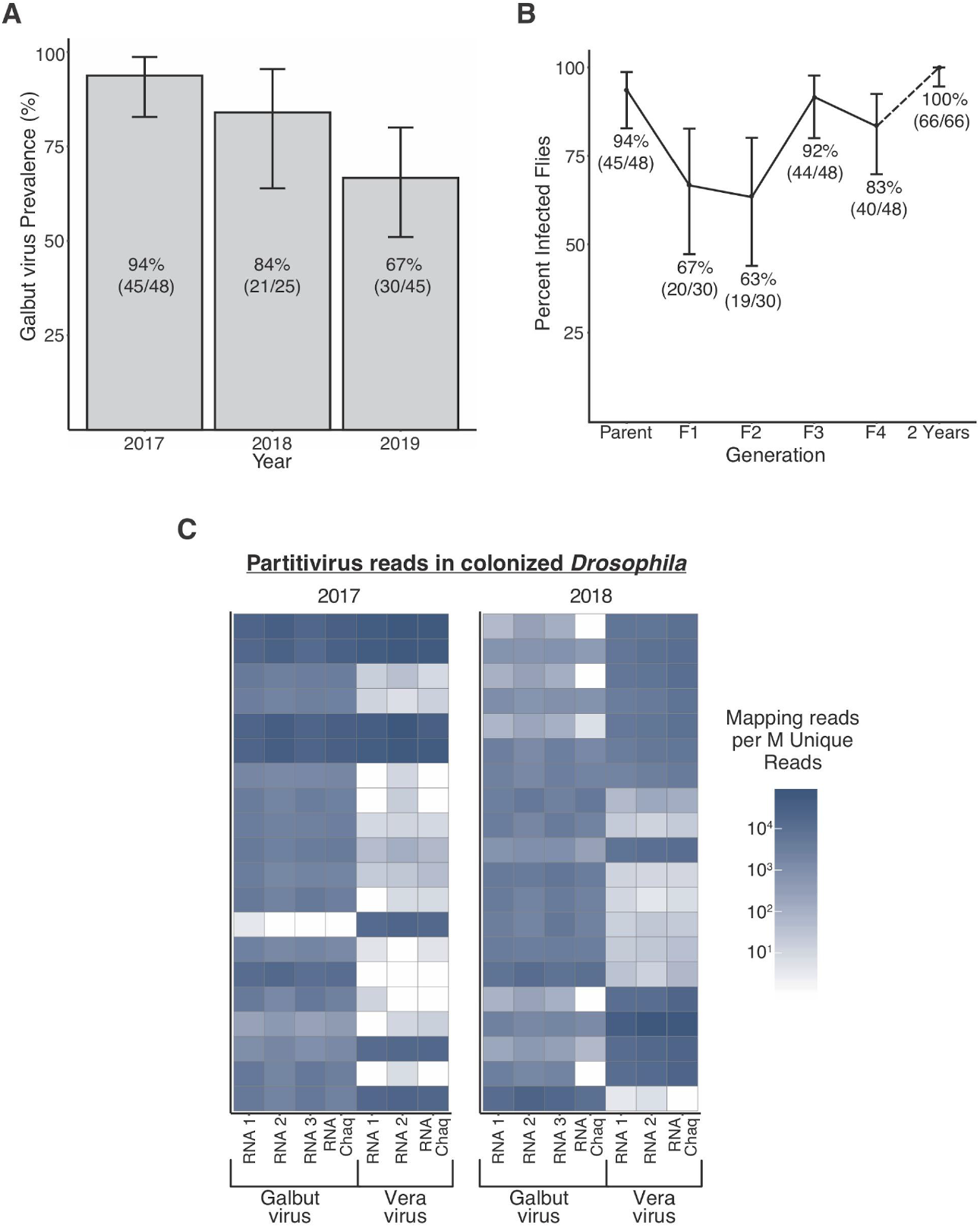
Galbut virus is found in wild populations consistently and can be maintained in colonized populations of *Drosophila melanogaster* over multiple years. A) Prevalence of galbut virus in wild-caught flies from the same backyard compost bin in Fort Collins, Colorado over 3 years. Numbers of flies positive by qRT-PCR and number of flies sampled are indicated. Error bars indicate 95% confidence intervals. B) Galbut virus was maintained for at least two years in colonized populations of wild *D. melanogaster* collected in Fort Collins, CO, USA in 2017 (FoCo-17). C) Metagenomic sequencing of individual *D. melanogaster* flies from colonies established in 2017 and 2018 shows prevalence and relative loads of galbut virus, vera virus, and their associated chaq and chaq-like segments. Each row represents an individual fly. Reads are normalized as mapping reads per million unique reads.

We first determined whether colonized flies would retain galbut virus infection, as a previous analysis of laboratory-reared *Drosophila* transcriptomic datasets showed a general absence of galbut virus (12). We tested galbut virus stability in the FoCo-17 population over four generations and then again after 2 years in the laboratory. We found that the parental generation had a prevalence of 93% while F1-F4 offspring maintained a prevalence between 63% and 92% (**Fig. 1B**). After two years, 66 out of 66 sampled FoCo-17 flies were infected (**Fig. 1B**). The FoCo-18 colony appears to have a similarly high prevalence of galbut virus infection months after establishment (**Fig. 1C**).

### Identification of additional partitiviruses in flies and mosquitoes

We identified the sequence of a second partitivirus in our metagenomic datasets from wild flies and from flies in our FoCo-17 and FoCo-18 colonies. We named this virus vera virus (vera means true in Esperanto). The vera virus genome consisted of an RNA encoding an RdRp (RNA #1), an RNA encoding a putative capsid protein (RNA #2), and a chaq virus-like RNA segment (**Fig. 2**). We did not identify any contigs with detectable sequence similarity to the predicted protein encoded on galbut virus RNA 3, nor contigs with similar occurrence patterns and coverage levels as vera virus RNAs 1 and 2. Like galbut virus, vera virus was detectable in wild-caught flies, and remained as a stable persistent infection in colonized populations over multiple years (**Fig. 1C**).

**Figure 2:**
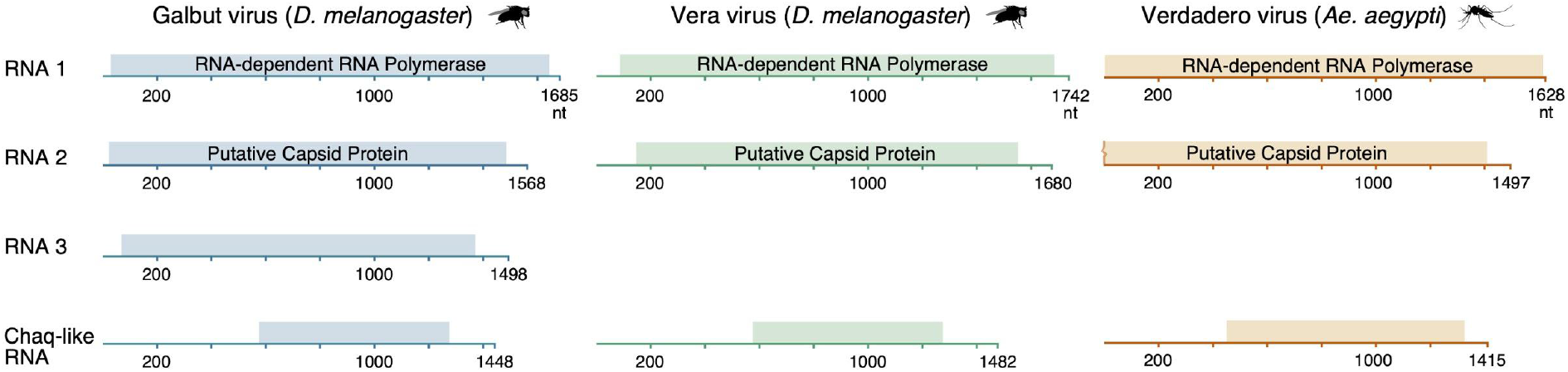
Genome structure of galbut virus, vera virus, and verdadero virus. Depictions of the genome structures of these viruses with predicted open reading frames indicated. RNA 1 of these viruses is predicted to encode RNA dependent RNA polymerases and RNA 2 a putative capsid protein. RNA 3 and the chaq virus-like segments are predicted to encode proteins of unknown function. The open reading frame on verdadero virus RNA 2 extends beyond the partial sequence that we recovered.

We also serendipitously identified another partitivirus as a persistent infection of a colony of *Aedes aegypti* mosquitoes originally derived from Poza Rica, Mexico. We named this virus verdadero virus (verdadero = true in Spanish, **Fig. 2**). The verdadero virus genome consisted of an RNA encoding an RdRp (RNA #1), an RNA encoding a putative capsid protein (RNA #2), and a chaq virus-like RNA segment. As was the case for vera virus, we could not identify a segment with similarity to RNA #3 of galbut virus. We tested 48 (24 male, 24 female) adult mosquitoes in the colony and 100% of these were positive for verdadero virus RNA by qRT-PCR [confidence interval 92.6-100%]. Other *Ae. aegypti* colonies housed at Colorado State University (strains New Orleans and Vergel) were negative for verdadero virus by qRT-PCR.

We created maximum likelihood phylogenies using these new partitivirus sequences and previously described similar sequences (**Fig. 3**). This tree includes a number of sequences from metagenomic surveys of invertebrates and some fungi-derived sequences. The invertebrate-derived sequences did not form a well-supported monophyletic cluster. In some cases, sequences from related hosts clustered together. For instance, Atrato Partiti-like virus 2 and Partitivirus-like 1 from *An. darlingi* and *An. gambiae* formed a well-supported group. But overall there was a notable lack of concordance by host. Mosquito-derived sequences were spread throughout the tree. Galbut virus and vera virus, both from the same populations of *Drosophila melanogaster*, were distantly placed on the tree.

**Figure 3:**
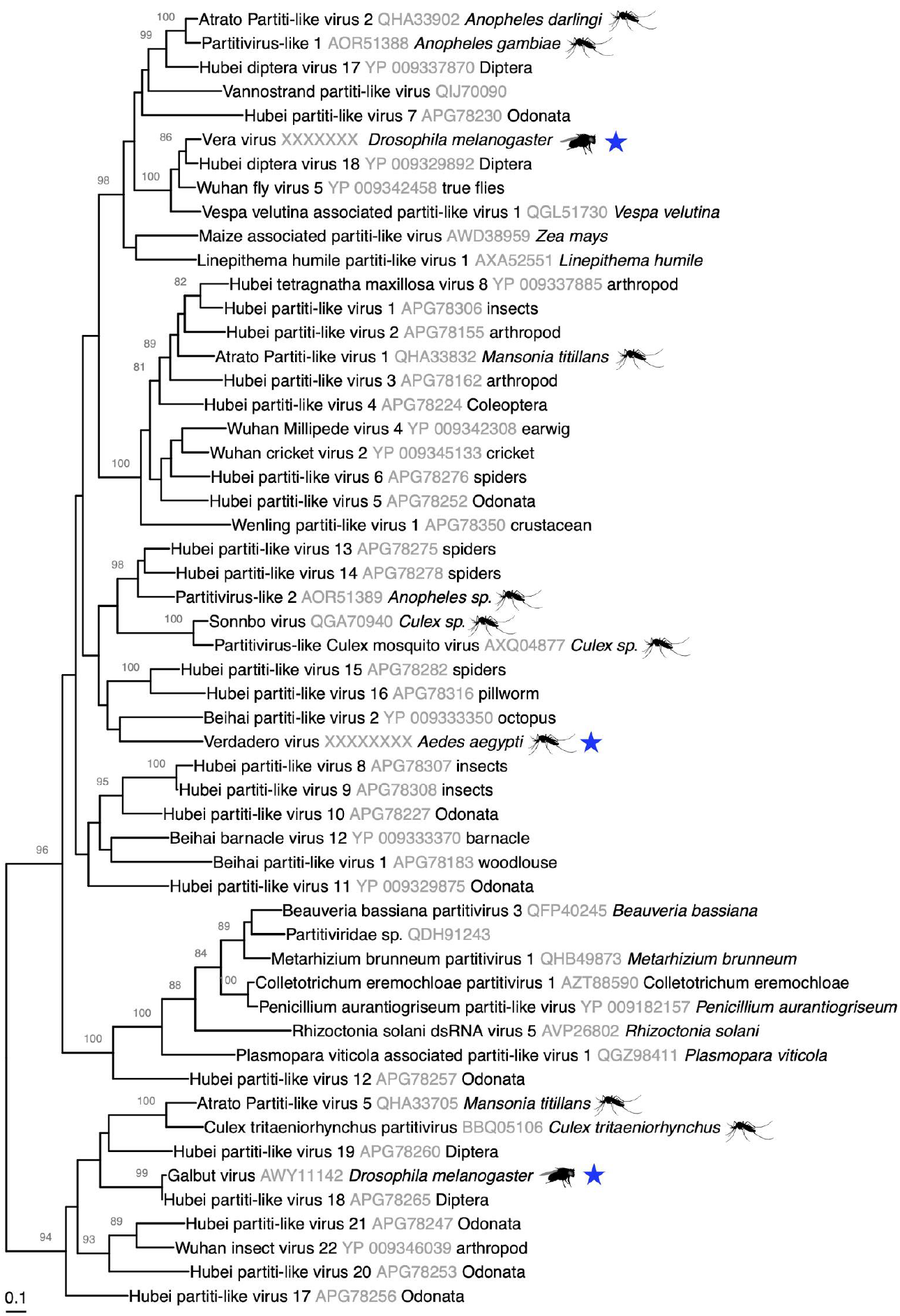
Phylogenetic tree of partitivirus-like sequences. Unrooted maximum likelihood phylogenetic tree of partitivirus-like RdRp sequences related to those described here (galbut virus, vera virus, and verdadero virus; marked with blue star). Sequences from mosquitoes and *Drosophila* are indicated with images. The host from each sequence’s metadata in NCBI is noted after the accession number. In many cases, exact hosts are uncertain because sequences derive from pools of different organisms.

### Galbut virus replicates in *Drosophila* cells

Two lines of evidence suggested that galbut virus actually infects flies. First, galbut virus derived small RNA profiles matched those expected for a virus that replicated in fly cells (12). Second, galbut virus clusters phylogenetically with a large group of partitivirus-like sequences from arthropod samples (**Fig. 3**). Nevertheless, direct evidence of galbut virus replication in fly cells had not been published, and it remained possible that galbut virus sequences actually derived from another microbial symbiont or were dietary in origin.

We therefore raised an antibody against the galbut virus putative capsid protein encoded on RNA segment 2. Laboratory strain w^*1118*^ flies tested negative for galbut virus RNA both by qRT-PCR and subsequent metagenomic sequencing. Extracts from FoCo-17 and w^*1118*^ flies were subjected to western blotting to confirm the specificity of the antibody. A band of the expected size, ~63 KDa, was present in galbut-infected FoCo-17 flies but not in uninfected w^*1118*^ flies (**Fig. 4A**). We used immunofluorescence microscopy using this antibody to identify sites of virus replication within sections of formalin fixed paraffin-embedded adult flies. We observed foci of staining within cells and tissues throughout adult male and female flies. Locations of infection included gut tissues (foregut, midgut, hindgut) and in egg chambers. These fluorescent foci were not evident in lab-reared, galbut virus uninfected w^*1118*^ flies (**Fig. 4B**).

**Figure 4:**
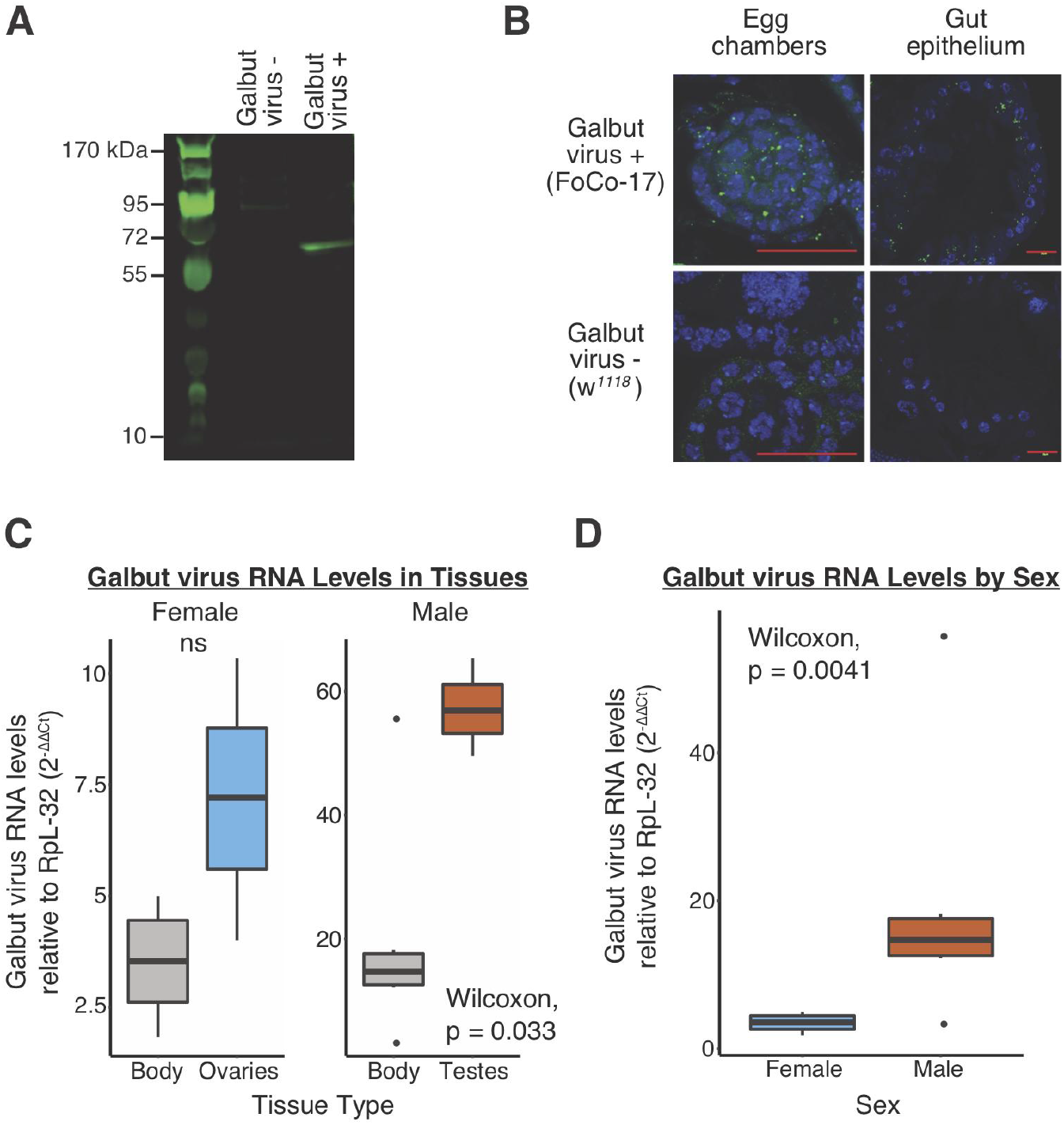
Screening of adult *Drosophila melanogaster* by immunofluorescence assay (IFA) and qPCR indicate galbut virus infects fly tissues. A) Galbut virus antibody specificity was confirmed by detection of a single band of the expected size of ~63kDa in infected flies (FoCo-17). B) Screening of galbut virus in adult FoCo-17 *D. melanogaster* by IFA showed foci of viral protein in cells in various tissues including the egg chambers and gut epithelium. Green color indicates staining by antibody raised against the putative galbut virus capsid protein; blue color indicates staining by Hoescht 33342 (DNA) C) Galbut virus RNA levels relative to the housekeeping gene RpL-32 (2^−ΔΔCt^ method) in dissected ovaries and testes as compared to adult bodies. Boxplots depict median values from 7 whole bodies of each sex and 3 pools per sex tissue type (10 testes or ovaries per pool). (Wilcoxon test, bonferroni adj., p = 0.033; ns: not significant). D) Galbut virus RNA levels relative to the housekeeping gene RpL-32 (2^−ΔΔCt^ method) in adult bodies compared by sex. Boxplots depict median values from 7 adult flies of each sex. (Wilcoxon test, bonferroni adj.; p = 0.0041).

Both ovaries and testes were positive for galbut virus RNA by qRT-PCR (**Fig. 4C**). In female flies, galbut virus RNA levels were higher in ovaries than RNA levels in whole bodies but not significantly so. Galbut virus RNA levels in testes were on average 3.9-fold higher than in whole male bodies (**Fig. 4C**; higher, Wilcoxon test, bonferroni adj., p = 0.033). When comparing galbut virus RNA levels in bodies by sex, we found RNA levels to be significantly higher (4.2-fold higher) in males than females (**Fig. 4D**; higher, Wilcoxon test, bonferroni adj., p = 0.0041). In general, galbut virus RNA levels were higher than those of ribosomal protein L32 (RpL32) mRNA, which is categorized as having “extremely high expression” in the modEncode database (51)

### Galbut virus exhibits efficient biparental vertical transmission

We set out to determine what modes of transmission could contribute to galbut virus’s exceptional high global prevalence. Having visualized viral protein in the egg chambers (**Fig. 4B**) and because dissected testes and ovaries tested positive for galbut virus by qRT-PCR (**Fig. 4C)**, we first evaluated maternal and paternal vertical transmission. We performed several experiments to quantify transmission efficiency. For all experiments, vertical transmission efficiency was quantified by testing for galbut virus RNA in adult offspring collected 2-5 days post eclosion.

In the first experiment, we set up individual crosses between FoCo-17 infected and w^*1118*^ uninfected adults, collected and bleached eggs, and placed eggs together in one vial per cross. Of the 6 independent maternal crosses (infected female, uninfected male), 34 of the 34 offspring were positive (100% transmission efficiency, confidence interval 89.7-100%; **Table 1**). Of the 5 independent paternal crosses (uninfected female, infected male), 27 of the 28 offspring tested positive (~96% transmission efficiency, confidence interval 81.7-99.9%; **Table 1**).

**Table 1:** *Drosophila melanogaster* and *Aedes aegypti* partitiviruses exhibit efficient biparental vertical transmission. Transmission efficiencies of vertical (maternal, paternal, and wild maternal) and horizontal (ingestion) routes. Numbers in brackets represent exact binomial confidence intervals.

| Experiment | Transmission | Species | Type | Group | # Crosses | # Screened* | Total Positive | Transmission Efficiency |
| --- | --- | --- | --- | --- | --- | --- | --- | --- |
| 1 | Vertical | *D. melanogaster* | Maternal | Female+/Male- | 6 | 34 | 34 | 100% [89.7, 100%] |
|  |  | *D. melanogaster* | Paternal | Female-/Male+ | 5 | 28 | 27 | 100% [81.7, 99.9%] |
| 2 | Vertical | *D. melanogaster* | Maternal | Female+/Male- | 4 | 44 | 44 | 100% [92.0, 100%] |
|  |  | *D. melanogaster* | Paternal | Female-/Male+ | 2 | 18 | 18 | 100% [81.5, 100%] |
| 3 | Vertical | *D. melanogaster* | Maternal | Wild Female+ | 7 | 120 | 119 | 99% [95.4, 100%] |
| 4 | Horizontal | *D. melanogaster* | Ingestion | Female | NA | 28 | 0 | 0% [0, 12.3%] |
|  |  | *D. melanogaster* | Ingestion | Male | NA | 26 | 0 | 0% [0, 13.2%] |
| 5 | Vertical | *Ae. aegypti* | Maternal | Female+/Male- | 4 | 48 | 48 | 100% [92.6, 100%] |
|  |  | *Ae. aegypti* | Paternal | Female-/Male+ | 4 | 39 | 38 | 97% [86.5, 99.9%] |

\*Offspring screened in vertical transmission, exposed adults screened in horizontal transmission

In the second experiment we separated bleached eggs into individual vials to avoid possible horizontal transmission between siblings during development. Of the 4 independent maternal crosses, 44 of 44 offspring tested positive (100% transmission efficiency, confidence interval 92.0-100%; **Table 1**). Of the 2 independent paternal crosses, 18 of 18 offspring tested positive (100% transmission efficiency, confidence interval 81.5-100%; **Table 1**).

In the third experiment, to verify that this high efficiency was not an artifact associated with laboratory-reared flies, we trapped wild female *D. melanogaster*, separated them into individual bottles, allowed them to lay eggs, and then tested mothers and offspring for galbut virus RNA. Offspring from 7 independent galbut virus positive females were tested, and we found a vertical transmission efficiency of 99% (119/120 offspring; confidence interval 95.4-100%; **Table 1**), indicating that efficient vertical transmission was not only associated with laboratory environments. We could not discern the infection status of the unknown fathers.

### No evidence of horizontal transmission by ingestion

To quantify galbut virus horizontal transmission (experiment 4), adult female and male flies were exposed through ingestion of homogenate derived from wild flies. This homogenate was infectious by microinjection (see below). Homogenate was mixed with a dyed food solution and placed in capillary feeding tubes. Ingestion was confirmed by visual inspection of fly abdomens. After feeding, flies were returned to normal food for 21 days. Following this incubation period, flies were collected and screened via qRT-PCR for galbut virus RNA. Virus levels in experimental flies were compared to virus loads in flies that were immediately flash frozen after ingestion of homogenate to establish an upper limit for residual RNA levels in the absence of viral replication. Although 3 of the 54 injected flies did test positive for galbut virus RNA after 21 days, their C_t_ values were on average 4.9 higher (~30 fold less RNA), than C_t_ values from immediately-frozen injected flies. We interpreted that these low level signals were likely derived from residual injected RNA. Therefore, we detected no evidence of active replication in any of the 54 flies tested (0% horizontal transmission; confidence interval 0-13.2%). This indicated that horizontal transmission, at least by ingestion, is unlikely to contribute substantially to maintenance of galbut virus infection in fly populations.

### A mosquito partitivirus also exhibits efficient biparental vertical transmission

We performed similar experiments to quantify vertical transmission efficiency of verdadero virus in *Aedes aegypti* by crossing infected female or male mosquitoes from the Poza Rica colony with uninfected counterparts from the New Orleans and Vergel colonies (experiment 5). Maternally, verdadero virus was transmitted at 100% efficiency (48/48 offspring, confidence interval 92.6-100%, **Table 1**) and paternally it was transmitted at 97% efficiency (38/39 offspring, confidence interval 86.5-99.9%, **Table 1**). This indicated that high vertical transmission efficiency is a characteristic of multiple dipteran infecting partitiviruses.

### Partitivirus RNA levels in *Drosophila* and mosquito offspring

Although both maternal and paternal transmission were ~100% efficient (**Table 1**), we were curious whether the exact route of vertical transmission impacted viral loads (total viral RNA relative to host reference genes) in offspring. In sigma virus, another biparentally transmitted virus in *Drosophila*, lower viral levels following paternal transmission lead to decreased paternal transmission in subsequent generations (52, 53). For galbut virus in *Drosophila*, offspring from paternal transmission (n=18) had significantly higher galbut virus RNA levels (5-fold median difference) compared to those infected by maternal transmission (n=44) (**Fig. 5A**, p = 0.0016). However, in *Aedes aegypti*, we saw the opposite trend: offspring from maternal transmission had significantly higher verdadero levels (107-fold median difference) than offspring infected via paternal transmission (**Fig. 5B**, p = 5.2×10^−6^). In both flies and mosquitoes, individual offspring exhibited broad distributions of viral loads that overlapped between sexes. Nevertheless, the route of vertical transmission impacted average partitivirus loads in offspring, and the direction of this effect was opposite for these two viruses.

**Figure 5:**
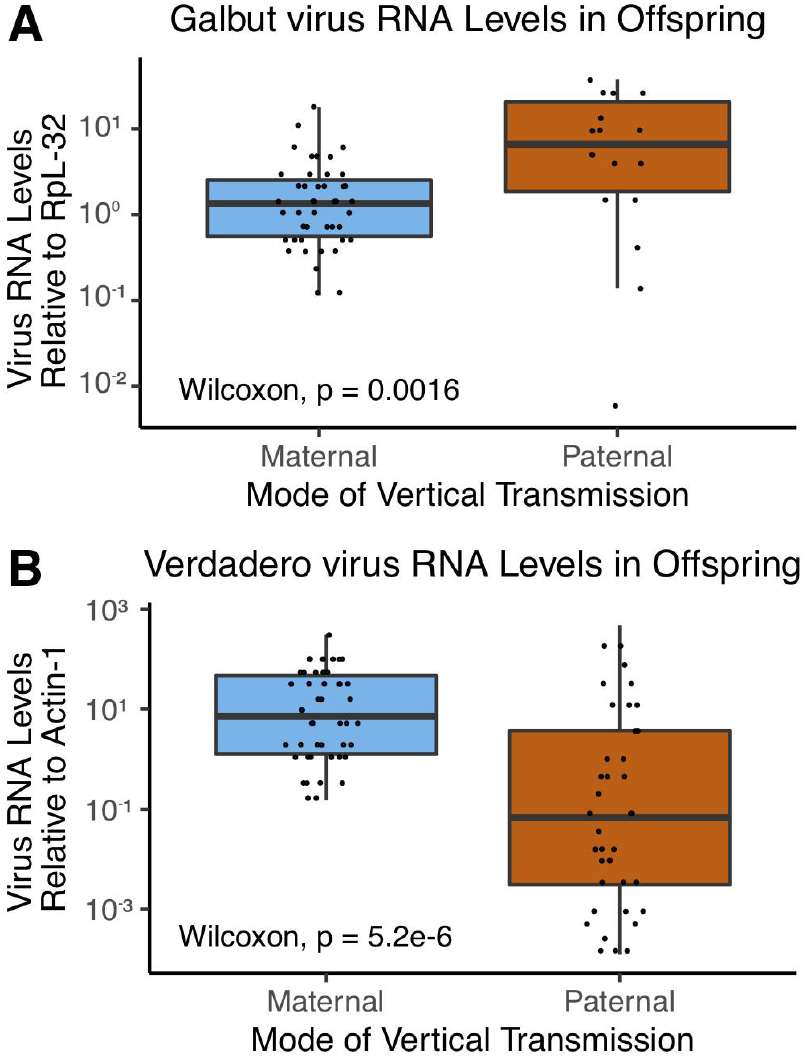
Galbut virus and verdadero virus RNA levels in offspring infected via maternal or paternal transmission. A) Galbut virus RNA levels in offspring (maternal n=44; paternal n=18) relative to the housekeeping gene RpL-32 (Wilcoxon test, p = 1.6×10^−3^). B) Verdadero virus RNA levels in offspring (maternal n=48; paternal n=38) relative to the housekeeping gene Actin-1 (Wilcoxon test, p = 5.2×10^−6^).

### Isolation of galbut virus through inbreeding

A challenge associated with the vast number of novel viruses being discovered by sequencing is their isolation for further biological characterization (5, 54). We attempted to isolate galbut virus in cell culture using *Drosophila* S2 cells but detected no evidence of replication. We also attempted to purify galbut virus particles by ultracentrifugation as described previously for Kallithea virus, but did not succeed, though this was not an exhaustive effort (55). We therefore designed a breeding scheme to create inbred fly lines that were only infected with galbut virus that took advantage of galbut virus’s efficient vertical transmission.

We randomly crossed pairs of virgins from the FoCo-17 population that harbored 5 viruses: galbut virus, vera virus, La Jolla virus, Nora virus, and Thika virus (12, 56). Eggs from 33 independent crosses were collected, bleached and placed in individual vials. This created a bottleneck to remove horizontally transmitted viruses and those with less efficient vertical transmission than galbut virus (57). Parents were retrospectively tested by qRT-PCR for all 5 viruses, and if both parents had fewer viruses than the previous generation, sibling offspring were randomly crossed again, eggs collected and screening repeated. We were able to generate a line detectable with only galbut virus after 4 generations, which we named FoCo-17 line #30 (**Fig. 6**). Also observed in this breeding scheme was the eventual purge of vera virus, the other partitivirus we identified in *Drosophila* (**Fig. 1, Fig. 2**). This suggested that although galbut virus and verdadero virus were transmitted efficiently at ~100% (**Table 1**), this efficiency may not be a universal feature of all arthropod-infecting partitiviruses.

**Figure 6:**
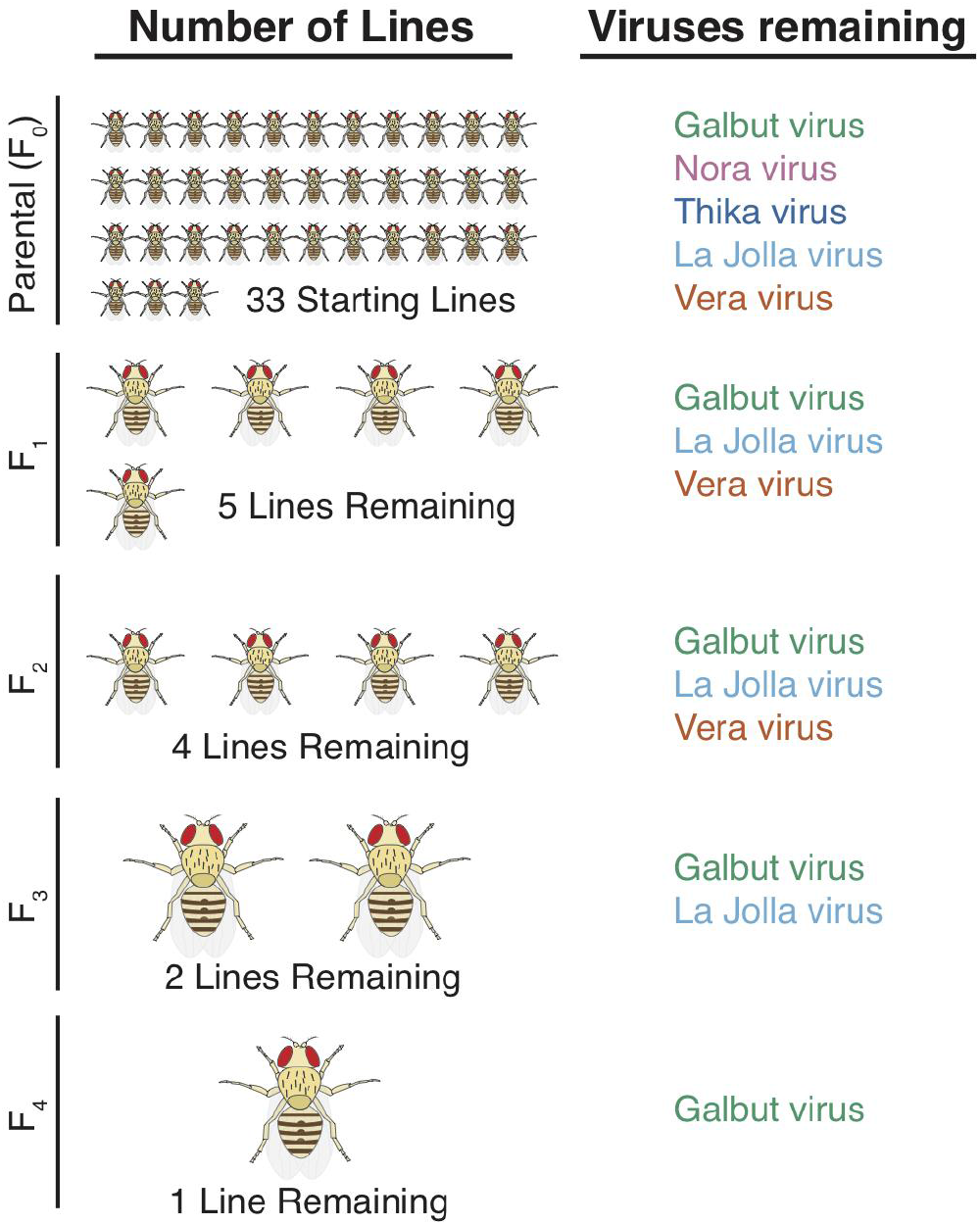
Creation of a line of *D. melanogaster* singly infected with galbut virus. Iterative inbreeding of *D. melanogaster* coupled to purging of lines that did not contain fewer viruses, lead to the establishment of a *D. melanogaster* line with galbut virus as the only detectable virus after four generations. Virus names on the right of the figure indicate the viruses present in any of the lines remaining at the indicated generation.

### Experimental infection

To establish a system for experimental infection we performed intrathoracic (IT) microinjection of virgin uninfected flies with a filtered homogenate from FoCo-17 line #30. We injected between 61 and 120 flies from each of three *Drosophila* Genetic Reference Panel (DGRP) strains (~50% of each sex) (34). 9 days post injection, we tested 5 male and 5 female injected flies per strain using qRT-PCR and in all cases 10/10 injected flies tested positive (**Table 2**). To verify that these flies were truly infected and that we were not detecting residual injected viral RNA, the remaining flies were crossed and we tested offspring from IT injected parents at 9, 15, and 19 days post injection. Offspring whose eggs were laid 9 or 15 days post injection were not uniformly infected (range 0-100%; **Table 2**). But, by 19 days post injection, all offspring tested were positive by qRT-PCR (**Table 2**). This confirmed that injected parents were legitimately infected and that infection had disseminated to tissues involved in vertical transmission in all DGRP strains by 15 days post injection. These infected offspring were used to establish colonies that differed from the original inbred DGRP lines only in their galbut virus infection status. Testing these colonies 50 days later (~4-5 generations), we found that DGRP strains 399 and 517 maintained 100% prevalence (24/24 flies for each positive by qRT-PCR, **Table 2**). However, DGRP 357 only had a galbut virus prevalence of ~21% (5/24 flies, **Table 2**). This suggested that multigenerational vertical transmission efficiency was a function of host genetic background.

**Table 2:**
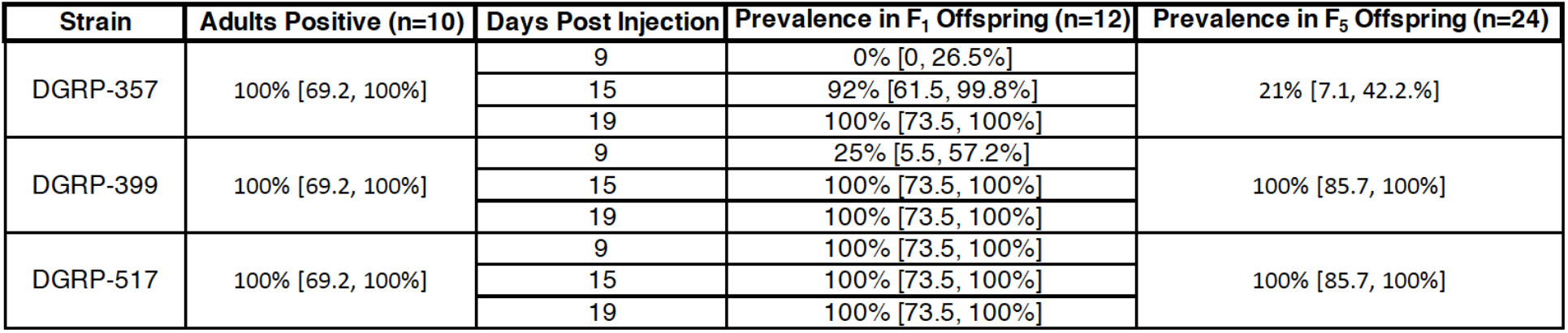
Experimental infection of galbut virus by microinjection and subsequent transmission to offspring. Adult flies from three DGRP strains were injected with a homogenate from galbut virus infected line #30. Ten injected flies per strain were tested for galbut virus RNA by qRT-PCR 9 days post injection; the percent positive of each set of 10 is indicated. Additional injected flies were allowed to mate, with subsequent testing of offspring from the first and 5th generation to monitor vertical transmission from injected parents and stability of transmission across multiple generations. First generation offspring whose eggs were laid 9, 15, or 19 days post injection were tested; the percent positive of each set of 12 flies are indicated. Infected colonies were established using offspring from days 15 and 19, and prevalence was tested 50 days after establishment (~4-5 generations). Numbers in brackets represent the lower and upper 95% confidence intervals. DGRP: *Drosophila* Genetic Reference Panel.

## Discussion

In this study, we began to characterize the biology of a recently discovered virus – galbut virus – that is unusually common in an important model organism. We confirmed that this “maybe” virus is indeed a virus of its putative fruit fly host and found that it can be transmitted efficiently from infected fathers or mothers to their offspring. This property is shared by at least one more insect-infecting partitivirus, verdadero virus, which we identified as a persistent infection in a colony of *Aedes aegypti* mosquitoes. This expands the known host range of the partitiviruses and suggests that the large number of partitivirus sequences that have been identified in a broad range of arthropods are likely legitimate viruses of those hosts. Efficient vertical transmission may be a mechanism that generally supports the success of these viruses.

An apparent paradox associated with this efficient biparental vertical transmission is that galbut virus has only been detected in ~60% of individual wild flies tested (**Fig. 1**, (12, 32)). Modeling indicates that highly efficient biparental transmission should eventually produce 100% of susceptible individuals to be infected, unless infection exacts a high fitness cost (58). Several hypotheses could account for this apparent discrepancy. First, it is possible that galbut virus is increasing or decreasing in frequency. In our sampling, prevalence decreased slightly over three years, though it is unclear whether this was just a stochastic effect nor whether these flies and viruses represented a single population lineage (**Fig. 1A**). Webster et al. suggested that the common ancestor of galbut virus in *D. melanogaster and D. simulans* populations existed ~200 years ago (12), and perhaps galbut virus has not setted to an equilibrium frequency. Additional longitudinal sampling will shine light on this. A related hypothesis is that galbut virus might exact enough of a fitness cost that it is driving an increase in host resistance alleles (59–62). This phenomenon has been observed for *D. melanogaster* sigmavirus: although sigma virus exhibits biparental vertical transmission, it negatively impacts host fitness resulting in prevalences of 0-30% worldwide (12, 63). Our experiments using different DGRP strains provided evidence that host genetics can modulate galbut virus transmission efficiency (**Table 2**). Contrary evidence is that we observed an increase in infection frequency to ~100% in 2 separately colonized populations (**Fig. 1C**). It may be that fitness costs are negligible in a laboratory environment. A final hypothesis is that variable communities of other microbiota in different individual flies could alter galbut virus infection efficiency (64, 65).

The efficient vertical transmission observed for these insect-infecting partitiviruses is reminiscent of what has been observed for their plant and fungus-infecting counterparts. Plant-infecting partitiviruses are completely dependent on vertical transmission, and horizontal transmission does not occur even when infected plants are grafted onto uninfected counterparts (19, 21, 24). Fungus-infecting partitiviruses also exhibit efficient vertical transmission (22–24), but the efficiency varies (66–72). Unlike the plant-infecting partitiviruses, fungus-infecting partitiviruses are able to transmit horizontally through processes such as hyphal anastomosis (22, 71, 73, 74). It appears that galbut virus and verdadero virus depend on vertical transmission like their plant-infecting counterparts (**Table 1**). Vera virus on the other hand, did not appear to have similarly efficient vertical transmission, as we were able to purge it relatively easily during the creation of *Drosophila* singly infected with galbut virus (**Fig. 6**).

Although we did not observe any evidence for horizontal transmission of galbut virus, there is evidence that galbut virus and other partitiviruses can transmit across species boundaries, so horizontal transmission does happen. First, there was a general lack of host-virus phylogenetic concordance, consistent with past cross-species transmission (**Fig. 3**). Second, galbut virus has been detected in a number of *Drosophila* species in the melanogaster group, including *D. simulans* and *D. suzukii* (12, 16, 32, 33). And, cross-species transmission of fungus-infecting partitiviruses has been documented (71, 75–77). Several possible mechanisms could allow cross-species transmission: The lack of horizontal transmission that we observed for galbut virus may not be representative of related viruses, and we only tested a single mode of horizontal infection. Other modes of horizontal transmission including sexual transmission, possibly during intraspecific mating attempts, could contribute. Mites or other parasites could serve as vectors for partitiviruses, as has been proposed for sigma viruses (78).

Some instances of cross species partitivirus transmission may involve jumps to distantly related organisms in other eukaryotic kingdoms. There is a well-supported clade of fungi-associated partitiviruses nested within the arthropod-associated partitiviruses (**Fig. 3)**. Similar phylogenetic interspersion of plant and fungus infecting partitiviruses has been noted (22, 79). Additional phylogenetic and experimental studies will be needed to address the intriguing possibility that partitiviruses are capable of long-range host switches.

Chaq virus was originally described as a virus-like sequence whose presence was correlated with galbut virus. It was postulated that chaq virus might be a satellite virus of galbut virus (chaq also means maybe, in Klingon) (12, 32). Consistent with previous reports, we found that chaq virus usually but not always co-occurred with galbut virus segments (**Fig. 1C**). We also identified chaq virus-like segments associated with vera and verdadero viruses, which are only distantly related to galbut virus (they share ~25-30% pairwise amino acid identity in their RdRp sequences). This indicates that if chaq-like segments are satellites of partitiviruses, this association has existed over long evolutionary time frames. An alternative interpretation is that chaq-like segments represent “optional” partitivirus segments not strictly required for replication, as has been described for multipartite mosquito-infecting jingmenviruses (80). A reverse genetics system for insect infecting partitiviruses would allow characterization of the function of individual viral proteins.

The highly efficient biparental vertical transmission that we documented for galbut and verdadero viruses is unusual for insect infecting viruses. Although many insect viruses are maintained both vertically and horizontally (81–84), there are several cases where insect viruses, like galbut virus, seem to be dependent on vertical transmission (63, 85, 86). In most documented cases however, maternal transmission is more efficient than paternal (63, 81, 86–88). An exception is rice stripe virus, where paternal transmission is more efficient in its leafhopper vector (89). The best precedent is probably *Drosophila*-infecting sigma viruses, which can be transmitted biparentally, allowing for sweeps through fly populations (63, 90). Parental transmission of *D. melanogaster* sigma virus is less efficient than maternal (~100% vs 65%), and paternally infected flies transmit infection less efficiently to the subsequent generation (52). We did observe different viral loads in offspring following paternal or maternal transmission, although for galbut virus, paternally infected flies had higher viral loads (**Fig. 5**). Whether a similar multigenerational phenomenon occurs for partitiviruses remains to be determined.

A particular challenge associated with understanding the biology of novel viruses identified via metagenomics is the inability to isolate them through classical methods such as cell culture. Many viruses are not culturable or not easily culturable. Here, we circumvent that obstacle through an iterative breeding scheme to generate flies with galbut virus as the only detectable infection (**Fig. 6**). We propose that this isolation method could be applied to other newly identified viruses. The isolation of galbut virus and the initial characterization described here establishes essential groundwork for further understanding the biological impacts and possible utility of arthropod-infecting partitiviruses.

## Acknowledgements

We thank Marylee Kapuscinski, Justin Lee, and the CSU Next Generation Sequencing Core Facility for assistance with sequencing, and Rebekah Kading for helpful discussion. We would like to acknowledge Dr. Bill Black for providing *Aedes aegypti* strains and Dr. Susan Tsunoda for providing *D. melanogaster* strain w^*1118*^.

## Funding

Stocks obtained from the Bloomington Drosophila Stock Center (NIH P40OD018537) were used in this study. This work was supported by the Colorado State University College of Veterinary Medicine and Biomedical Sciences College Research Council. Computational resources were supported by NIH/NCATS Colorado CTSA Grant Number UL1 TR002535. STC was supported in part by National Science Foundation (NSF) NRT grant 1450032. MRR was supported as a NSF Graduate Fellow (Grant No. 5325000). BDF was supported by NIH/NIAID grants R21AI129464 and U01AI138910. Any opinions, findings, conclusions or recommendations expressed in this paper are those of the author(s) and do not necessarily reflect the views of the funding organizations.

## References cited

1. Li C-X, Shi M, Tian J-H, Lin X-D, Kang Y-J, Chen L-J, Qin X-C, Xu J, Holmes EC, Zhang Y-Z. 2015. Unprecedented genomic diversity of RNA viruses in arthropods reveals the ancestry of negative-sense RNA viruses. eLife 4:e05378.

2. Shi M, Lin X-D, Tian J-H, Chen L-J, Chen X, Li C-X, Qin X-C, Li J, Cao J-P, Eden J-S, Buchmann J, Wang W, Xu J, Holmes EC, Zhang Y-Z. 2016. Redefining the invertebrate RNA virosphere. Nature 540:539.

3. Wang D. 2015. Fruits of Virus Discovery: New Pathogens and New Experimental Models. Journal of Virology 89:1486–1488.

4. Tisza MJ, Pastrana DV, Welch NL, Stewart B, Peretti A, Starrett GJ, Pang Y-YS, Krishnamurthy SR, Pesavento PA, McDermott DH, Murphy PM, Whited JL, Miller B, Brenchley J, Rosshart SP, Rehermann B, Doorbar J, Ta’ala BA, Pletnikova O, Troncoso JC, Resnick SM, Bolduc B, Sullivan MB, Varsani A, Segall AM, Buck CB. 2020. Discovery of several thousand highly diverse circular DNA viruses. Elife 9.

5. Greninger AL. 2018. A decade of RNA virus metagenomics is (not) enough. Virus Res 244:218–229.

6. Brum JR, Ignacio-Espinoza JC, Roux S, Doulcier G, Acinas SG, Alberti A, Chaffron S, Cruaud C, Vargas C de, Gasol JM, Gorsky G, Gregory AC, Guidi L, Hingamp P, Iudicone D, Not F, Ogata H, Pesant S, Poulos BT, Schwenck SM, Speich S, Dimier C, Kandels-Lewis S, Picheral M, Searson S, Coordinators TO, Bork P, Bowler C, Sunagawa S, Wincker P, Karsenti E, Sullivan MB. 2015. Patterns and ecological drivers of ocean viral communities. Science 348.

7. Yutin N, Bäckström D, Ettema TJG, Krupovic M, Koonin EV. 2018. Vast diversity of prokaryotic virus genomes encoding double jelly-roll major capsid proteins uncovered by genomic and metagenomic sequence analysis. Virol J 15:67.

8. Sadeghi M, Altan E, Deng X, Barker CM, Fang Y, Coffey LL, Delwart E. 2018. Virome of > 12 thousand Culex mosquitoes from throughout California. Virology 523:74–88.

9. Roossinck MJ, Martin DP, Roumagnac P. 2015. Plant Virus Metagenomics: Advances in Virus Discovery. Phytopathology® 105:716–727.

10. Shi M, Lin X-D, Chen X, Tian J-H, Chen L-J, Li K, Wang W, Eden J-S, Shen J-J, Liu L, Holmes EC, Zhang Y-Z. 2018. The evolutionary history of vertebrate RNA viruses. Nature 556:197–202.

11. Koonin EV, Dolja VV. 2018. Metaviromics: a tectonic shift in understanding virus evolution. Virus Research 246:A1–A3.

12. Webster CL, Waldron FM, Robertson S, Crowson D, Ferrari G, Quintana JF, Brouqui J-M, Bayne EH, Longdon B, Buck AH, Lazzaro BP, Akorli J, Haddrill PR, Obbard DJ. 2015. The Discovery, Distribution, and Evolution of Viruses Associated with Drosophila melanogaster. PLOS Biology 13:e1002210.

13. Coyle MC, Elya CN, Bronski MJ, Eisen MB. 2018. Entomophthovirus: An insect-derived iflavirus that infects a behavior manipulating fungal pathogen of dipterans.

14. Fauver JR, Grubaugh ND, Krajacich BJ, Weger-Lucarelli J, Lakin SM, Fakoli LS, Bolay FK, Diclaro JW, Dabiré KR, Foy BD, Brackney DE, Ebel GD, Stenglein MD. 2016. West African Anopheles gambiae mosquitoes harbor a taxonomically diverse virome including new insect-specific flaviviruses, mononegaviruses, and totiviruses. Virology 498:288–299.

15. Li C-X, Shi M, Tian J-H, Lin X-D, Kang Y-J, Chen L-J, Qin X-C, Xu J, Holmes EC, Zhang Y-Z. 2015. Unprecedented genomic diversity of RNA viruses in arthropods reveals the ancestry of negative-sense RNA viruses. Elife 4.

16. Medd NC, Fellous S, Waldron FM, Xuéreb A, Nakai M, Cross JV, Obbard DJ. 2018. The virome of Drosophila suzukii, an invasive pest of soft fruit. Virus Evolution 4.

17. Lara Pinto AZ de, Santos de Carvalho M, de Melo FL, Ribeiro ALM, Morais Ribeiro B, Dezengrini Slhessarenko R. 2017. Novel viruses in salivary glands of mosquitoes from sylvatic Cerrado, Midwestern Brazil. PLOS ONE 12:e0187429.

18. Faizah AN, Kobayashi D, Isawa H, Amoa-Bosompem M, Murota K, Higa Y, Futami K, Shimada S, Kim KS, Itokawa K, Watanabe M, Tsuda Y, Minakawa N, Miura K, Hirayama K, Sawabe K. 2020. Deciphering the Virome of Culex vishnui Subgroup Mosquitoes, the Major Vectors of Japanese Encephalitis, in Japan. Viruses 12:264.

19. Boccardo G, Lisa V, Luisoni E, Milne RG. 1987. Cryptic plant viruses. Adv Virus Res 32:171–214.

20. Ghabrial S, Ochoa W, Baker T, Nibert M. 2008. Partitiviruses: General Features. Encyclopedia of Virology 68–75.

21. Roossinck MJ. 2010. Lifestyles of plant viruses. Philos Trans R Soc Lond, B, Biol Sci 365:1899–1905.

22. Nibert ML, Ghabrial SA, Maiss E, Lesker T, Vainio EJ, Jiang D, Suzuki N. 2014. Taxonomic reorganization of family Partitiviridae and other recent progress in partitivirus research. Virus Research 188:128–141.

23. Ghabrial SA, Castón JR, Jiang D, Nibert ML, Suzuki N. 2015. 50-plus years of fungal viruses. Virology 479–480:356–368.

24. Vainio EJ, Chiba S, Ghabrial SA, Maiss E, Roossinck M, Sabanadzovic S, Suzuki N, Xie J, Nibert M, ICTV Report Consortium. 2018. ICTV Virus Taxonomy Profile: Partitiviridae. Journal of General Virology, 99:17–18.

25. Jenkins MC, Higgins J, Abrahante JE, Kniel KE, BO’rien C, Trout J, Lancto CA, Abrahamsen MS, Fayer R. 2008. Fecundity of Cryptosporidium parvum is correlated with intracellular levels of the viral symbiont CPV. International Journal for Parasitology 38:1051–1055.

26. Xiao X, Cheng J, Tang J, Fu Y, Jiang D, Baker TS, Ghabrial SA, Xie J. 2014. A Novel Partitivirus That Confers Hypovirulence on Plant Pathogenic Fungi. Journal of Virology 88:10120–10133.

27. Zheng L, Zhang M, Chen Q, Zhu M, Zhou E. 2014. A novel mycovirus closely related to viruses in the genus Alphapartitivirus confers hypovirulence in the phytopathogenic fungus Rhizoctonia solani. Virology 456–457:220–226.

28. Sasaki A, Nakamura H, Suzuki N, Kanematsu S. 2016. Characterization of a new megabirnavirus that confers hypovirulence with the aid of a co-infecting partitivirus to the host fungus, Rosellinia necatrix. Virus Research 219:73–82.

29. Kamaruzzaman M, He G, Wu M, Zhang J, Yang L, Chen W, Li G. 2019. A Novel Partitivirus in the Hypovirulent Isolate QT5-19 of the Plant Pathogenic Fungus Botrytis cinerea. Viruses 11:24.

30. Safari M, Ferrari MJ, Roossinck MJ. 2019. Manipulation of Aphid Behavior by a Persistent Plant Virus. J Virol 93:e01781–18.

31. Pettersson JH-O, Shi M, Eden J-S, Holmes EC, Hesson JC. 2019. Meta-Transcriptomic Comparison of the RNA Viromes of the Mosquito Vectors Culex pipiens and Culex torrentium in Northern Europe. 11. Viruses 11:1033.

32. Shi M, White VL, Schlub T, Eden J-S, Hoffmann AA, Holmes EC. 2018. No detectable effect of *Wolbachia w* Mel on the prevalence and abundance of the RNA virome of *Drosophila melanogaster*. Proceedings of the Royal Society B: Biological Sciences 285:20181165.

33. Webster CL, Longdon B, Lewis SH, Obbard DJ. 2016. Twenty-Five New Viruses Associated with the Drosophilidae (Diptera). Evol Bioinform Online 12:13–25.

34. Mackay TFC, Richards S, Stone EA, Barbadilla A, Ayroles JF, Zhu D, Casillas S, Han Y, Magwire MM, Cridland JM, Richardson MF, Anholt RRH, Barrón M, Bess C, Blankenburg KP, Carbone MA, Castellano D, Chaboub L, Duncan L, Harris Z, Javaid M, Jayaseelan JC, Jhangiani SN, Jordan KW, Lara F, Lawrence F, Lee SL, Librado P, Linheiro RS, Lyman RF, Mackey AJ, Munidasa M, Muzny DM, Nazareth L, Newsham I, Perales L, Pu L-L, Qu C, Ràmia M, Reid JG, Rollmann SM, Rozas J, Saada N, Turlapati L, Worley KC, Wu Y-Q, Yamamoto A, Zhu Y, Bergman CM, Thornton KR, Mittelman D, Gibbs RA. 2012. The Drosophila melanogaster Genetic Reference Panel. Nature 482:173–178.

35. Vera-Maloof FZ, Saavedra-Rodriguez K, Elizondo-Quiroga AE, Lozano-Fuentes S, Black Iv WC. 2015. Coevolution of the Ile1,016 and Cys1,534 Mutations in the Voltage Gated Sodium Channel Gene of Aedes aegypti in Mexico. PLoS Negl Trop Dis 9:e0004263–e0004263.

36. Steven Denham, Lars Eisen, Meaghan Beaty, Barry J. Beaty, William C. Black, Karla Saavedra-Rodriguez. 2015. Two Novel Bioassays to Assess the Effects of Pyrethroid-Treated Netting on Knockdown-Susceptible Versus Resistant Strains of Aedes aegypti. Journal of the American Mosquito Control Association 31:52–62.

37. Magalhaes T, Bergren NA, Bennett SL, Borland EM, Hartman DA, Lymperopoulos K, Sayre R, Borlee BR, Campbell CL, Foy BD, Olson KE, Blair CD, Black W, Kading RC. 2019. Induction of RNA interference to block Zika virus replication and transmission in the mosquito Aedes aegypti. Insect Biochemistry and Molecular Biology 111:103169.

38. Clopper CJ, Pearson ES. 1934. THE USE OF CONFIDENCE OR FIDUCIAL LIMITS ILLUSTRATED IN THE CASE OF THE BINOMIAL. Biometrika 26:404–413.

39. Zer S, Ryvkin J, Wilner HJ, Zak H, Shmueli A, Shohat-Ophir G. 2016. A Simple Way to Measure Alterations in Reward-seeking Behavior Using Drosophila melanogaster. J Vis Exp 54910.

40. Cross S, Kapuscinski M, Perino J, Maertens B, Weger-Lucarelli J, Ebel G, Stenglein M. 2018. Co-Infection Patterns in Individual Ixodes scapularis Ticks Reveal Associations between Viral, Eukaryotic and Bacterial Microorganisms. Viruses 10:388.

41. Altschul SF, Gish W, Miller W, Myers EW, Lipman DJ. 1990. Basic local alignment search tool. J Mol Biol 215:403–410.

42. Li W, Godzik A. 2006. Cd-hit: a fast program for clustering and comparing large sets of protein or nucleotide sequences. Bioinformatics 22:1658–1659.

43. Langmead B, Salzberg SL. 2012. Fast gapped-read alignment with Bowtie 2. Nat Methods 9:357–359.

44. Katoh K, Standley DM. 2013. MAFFT Multiple Sequence Alignment Software Version 7: Improvements in Performance and Usability. Mol Biol Evol 30:772–780.

45. Capella-Gutiérrez S, Silla-Martínez JM, Gabaldón T. 2009. trimAl: a tool for automated alignment trimming in large-scale phylogenetic analyses. Bioinformatics 25:1972–1973.

46. Darriba D, Posada D, Kozlov AM, Stamatakis A, Morel B, Flouri T. 2020. ModelTest-NG: A New and Scalable Tool for the Selection of DNA and Protein Evolutionary Models. Mol Biol Evol 37:291–294.

47. Kozlov AM, Darriba D, Flouri T, Morel B, Stamatakis A. 2019. RAxML-NG: a fast, scalable and user-friendly tool for maximum likelihood phylogenetic inference. Bioinformatics 35:4453–4455.

48. Yu G, Smith DK, Zhu H, Guan Y, Lam TT-Y. 2017. ggtree: an r package for visualization and annotation of phylogenetic trees with their covariates and other associated data. Methods in Ecology and Evolution 8:28–36.

49. Emery P. 2007. Protein Extraction From Drosophila Heads, p. 375–377. *In* Rosato, E (ed.), Circadian Rhythms: Methods and Protocols. Humana Press, Totowa, NJ.

50. Stenglein MD, Sanders C, Kistler AL, Ruby JG, Franco JY, Reavill DR, Dunker F, DeRisi JL. 2012. Identification, Characterization, and In Vitro Culture of Highly Divergent Arenaviruses from Boa Constrictors and Annulated Tree Boas: Candidate Etiological Agents for Snake Inclusion Body Disease. mBio 3.

51. Celniker SE, Dillon LAL, Gerstein MB, Gunsalus KC, Henikoff S, Karpen GH, Kellis M, Lai EC, Lieb JD, MacAlpine DM, Micklem G, Piano F, Snyder M, Stein L, White KP, Waterston RH, modENCODE Consortium. 2009. Unlocking the secrets of the genome. Nature 459:927–930.

52. Fleuriet A. 1988. Maintenance of a Hereditary Virus, p. 1–30. *In* Hecht, MK, Wallace, B (eds.), Evolutionary Biology: Volume 23. Springer US, Boston, MA.

53. Longdon B, Wilfert L, Obbard DJ, Jiggins FM. 2011. Rhabdoviruses in two species of Drosophila: vertical transmission and a recent sweep. Genetics 188:141–150.

54. Wang D. 2020. 5 challenges in understanding the role of the virome in health and disease. PLOS Pathogens 16:e1008318.

55. Palmer WH, Medd NC, Beard PM, Obbard DJ. 2018. Isolation of a natural DNA virus of Drosophila melanogaster, and characterisation of host resistance and immune responses. PLoS Pathog 14:e1007050.

56. Habayeb MS, Ekengren SK, Hultmark D. 2006. Nora virus, a persistent virus in Drosophila, defines a new picorna-like virus family. J Gen Virol 87:3045–3051.

57. Merkling SH, van Rij RP. 2015. Analysis of resistance and tolerance to virus infection in Drosophila. Nat Protoc 10:1084–1097.

58. Fine PE. 1975. Vectors and vertical transmission: an epidemiologic perspective. Ann N Y Acad Sci 266:173–194.

59. Magwire MM, Fabian DK, Schweyen H, Cao C, Longdon B, Bayer F, Jiggins FM. 2012. Genome-wide association studies reveal a simple genetic basis of resistance to naturally coevolving viruses in Drosophila melanogaster. PLoS Genet 8:e1003057.

60. Xu J, Cherry S. 2014. Viruses and Antiviral Immunity in Drosophila. Dev Comp Immunol 42.

61. Swevers L, Liu J, Smagghe G. 2018. Defense Mechanisms against Viral Infection in Drosophila: RNAi and Non-RNAi. Viruses 10.

62. Palmer WH, Varghese FS, van Rij RP. 2018. Natural Variation in Resistance to Virus Infection in Dipteran Insects. Viruses 10.

63. Longdon B, Jiggins FM. 2012. Vertically transmitted viral endosymbionts of insects: do sigma viruses walk alone? Proc Biol Sci 279:3889–3898.

64. Sansone CL, Cohen J, Yasunaga A, Xu J, Osborn G, Subramanian H, Gold B, Buchon N, Cherry S. 2015. Microbiota-Dependent Priming of Antiviral Intestinal Immunity in Drosophila. Cell Host Microbe 18:571–581.

65. Faria VG, Martins NE, Magalhães S, Paulo TF, Nolte V, Schlötterer C, Sucena É, Teixeira L. 2016. Drosophila Adaptation to Viral Infection through Defensive Symbiont Evolution. PLOS Genetics 12:e1006297.

66. Ghabrial SA. 1998. Origin, Adaptation and Evolutionary Pathways of Fungal Viruses. Virus Genes 16:119–131.

67. Varga J, Rinyu E, Kevei É, Tóth B, Kozakiewicz Z. 1998. Double-stranded RNA mycoviruses in species of Aspergillus sections Circumdati and Fumigati. Can J Microbiol 44:569–574.

68. Anagnostakis SL, Chen B, Geletka LM, Nuss DL. 1998. Hypovirus Transmission to Ascospore Progeny by Field-Released Transgenic Hypovirulent Strains of Cryphonectria parasitica. Phytopathology® 88:598–604.

69. Coenen A, Kevei F, Hoekstra RF. 1997. Factors affecting the spread of double-stranded RNA viruses in Aspergillus nidulans. Genetical Research1997/02/01. 69:1–10.

70. Chun SJ, Lee Y-H. 1997. Inheritance of dsRNAs in the rice blast fungus, Magnaporthe grisea. FEMS Microbiology Letters 148:159–162.

71. Ihrmark K, Johannesson H, Stenström E, Stenlid J. 2002. Transmission of double-stranded RNA in Heterobasidion annosum. Fungal Genetics and Biology 36:147–154.

72. Ihrmark K, Stenström E, Stenlid J. 2004. Double-stranded RNA transmission through basidiospores of Heterobasidion annosum. Mycological Research 108:149–153.

73. Sasaki A, Kanematsu S, Onoue M, Oyama Y, Yoshida K. 2006. Infection of Rosellinia necatrix with purified viral particles of a member of Partitiviridae (RnPV1-W8). Archives of Virology 151:697–707.

74. Kanematsu S, Sasaki A, Onoue M, Oikawa Y, Ito T. 2010. Extending the Fungal Host Range of a Partitivirus and a Mycoreovirus from Rosellinia necatrix by Inoculation of Protoplasts with Virus Particles. Phytopathology® 100:922–930.

75. Vainio EJ, Hakanpää J, Dai Y-C, Hansen E, Korhonen K, Hantula J. 2011. Species of Heterobasidion host a diverse pool of partitiviruses with global distribution and interspecies transmission. Fungal Biology 115:1234–1243.

76. Vainio EJ, Korhonen K, Tuomivirta TT, Hantula J. 2010. A novel putative partitivirus of the saprotrophic fungus Heterobasidion ecrustosum infects pathogenic species of the Heterobasidion annosum complex. Fungal Biology 114:955–965.

77. Filippou C, Garrido-Jurado I, Meyling VN, Quesada-Moraga E, Coutts HAR, Kotta-Loizou I. 2018. Mycoviral Population Dynamics in Spanish Isolates of the Entomopathogenic Fungus Beauveria bassiana. Viruses 10.

78. Longdon B, Obbard DJ, Jiggins FM. 2010. Sigma viruses from three species of Drosophila form a major new clade in the rhabdovirus phylogeny. Proc Biol Sci 277:35–44.

79. Szegő A, Enünlü N, Deshmukh SD, Veliceasa D, Hunyadi-Gulyás É, Kühne T, Ilyés P, Potyondi L, Medzihradszky K, Lukács N. 2010. The genome of Beet cryptic virus 1 shows high homology to certain cryptoviruses present in phylogenetically distant hosts. Virus Genes 40:267–276.

80. Ladner JT, Wiley MR, Beitzel B, Auguste AJ, Dupuis AP, Lindquist ME, Sibley SD, Kota KP, Fetterer D, Eastwood G, Kimmel D, Prieto K, Guzman H, Aliota MT, Reyes D, Brueggemann EE, John LSt, Hyeroba D, Lauck M, Friedrich TC, O’Connor DH, Gestole MC, Cazares LH, Popov VL, Castro-Llanos F, Kochel TJ, Kenny T, White B, Ward MD, Loaiza JR, Goldberg TL, Weaver SC, Kramer LD, Tesh RB, Palacios G. 2016. A multicomponent animal virus isolated from mosquitoes. Cell Host Microbe 20:357–367.

81. Williams T, Virto C, Murillo R, Caballero P. 2017. Covert Infection of Insects by Baculoviruses. Frontiers in Microbiology 8:1337.

82. Agboli E, Leggewie M, Altinli M, Schnettler E. 2019. Mosquito-Specific Viruses-Transmission and Interaction. Viruses 11:873.

83. Chen Y, Evans J, Feldlaufer M. 2006. Horizontal and vertical transmission of viruses in the honey bee, Apis mellifera. Journal of Invertebrate Pathology 92:152–159.

84. Nakashima N, Noda H. 1995. Nonpathogenic Nilaparvata lugens Reovirus Is Transmitted to the Brown Planthopper through Rice Plant. Virology 207:303–307.

85. Strand MR, Burke GR. 2013. Polydnavirus-wasp associations: evolution, genome organization, and function. Current Opinion in Virology 3:587–594.

86. Chen Q, Godfrey K, Liu J, Mao Q, Kuo Y-W, Falk BW. 2019. A Nonstructural Protein Responsible for Viral Spread of a Novel Insect Reovirus Provides a Safe Channel for Biparental Virus Transmission to Progeny. J Virol 93:e00702–19.

87. Ferber ML, Rı́os AF, Kuhl G, Comendador MA, Louis C. 1997. Infection of the Gonads of the SimES Strain ofDrosophila simulansby the Hereditary Reovirus DSV. Journal of Invertebrate Pathology 70:143–149.

88. Longdon B, Day JP, Schulz N, Leftwich PT, de Jong MA, Breuker CJ, Gibbs M, Obbard DJ, Wilfert L, Smith SCL, McGonigle JE, Houslay TM, Wright LI, Livraghi L, Evans LC, Friend LA, Chapman T, Vontas J, Kambouraki N, Jiggins FM. Vertically transmitted rhabdoviruses are found across three insect families and have dynamic interactions with their hosts 9.

89. Mao Q, Wu W, Liao Z, Li J, Jia D, Zhang X, Chen Q, Chen H, Wei J, Wei T. 2019. Viral pathogens hitchhike with insect sperm for paternal transmission. Nature Communications 10:955.

90. Longdon B, Wilfert L, Obbard DJ, Jiggins FM. 2011. Rhabdoviruses in Two Species of Drosophila: Vertical Transmission and a Recent Sweep. Genetics 188:141.

